# MoMo: Discovery of statistically significant post-translational modification motifs

**DOI:** 10.1101/410050

**Authors:** Alice Cheng, Charles E. Grant, William S. Noble, Timothy L. Bailey

**Affiliations:** Department of Genome Sciences, University of Washington, Seattle, Washington, USA; Department of Computer Science and Engineering, University of Washington, Seattle, Washington, USA; Department of Pharmacology, University of Nevada, Reno, Nevada, 89557, USA

## Abstract

**Motivation:** Post-translational modifications (PTMs) of proteins are associated with many significant biological functions and can be identified in high throughput using tandem mass spectrometry. Many PTMs are associated with short sequence patterns called “motifs” that help localize the modifying enzyme. Accordingly, many algorithms have been designed to identify these motifs from mass spectrometry data. Accurate statistical confidence estimates for discovered motifs are critically important for proper interpretation and in the design of downstream experimental validation.

**Results:** We describe a method for assigning statistical confidence estimates to PTM motifs, and we demonstrate that this method provides accurate *p*-values on both simulated and real data. Our methods are implemented in MoMo, a software tool for discovering motifs among sets of PTMs that we make available as a web server and as downloadable source code. MoMo reimplements the two most widely used PTM motif discovery algorithms—motif-x and MoDL—while offering many enhancements. Relative to motif-x, MoMo offers improved statistical confidence estimates and more accurate calculation of motif scores. The MoMo web server offers more proteome databases, more input formats, larger inputs and longer running times than the motif-x web server. Finally, our study demonstrates that the confidence estimates produced by motif-x are inaccurate. This inaccuracy stems in part from the common practice of drawing “background” peptides from an unshuffled proteome database. Our results thus suggest that many of the hundreds of papers that use motif-x to find motifs may be reporting results that lack statistical support.

**Availability:** http://meme-suite.org

## 1 Introduction

Protein post-translational modifications control or otherwise participate in a wide range of biological activities, including critical regulatory functions. Dysregulation of some types of modifications, such as phosphorylation, has been implicated in a variety of diseases, most notably cancer [8]. The best characterized modifications include methylation, acetylation, phosphorylation, and sumoylation, but hundreds of less common types of modifications have been identified and cataloged in various databases, including 1447 in UniMod. Additionally, thousands of individual modification sites have been identified experimentally in diverse organisms and are cataloged in generic PTM databases [9, 16] and databases devoted to specific types of PTMs [6, 4, 8, 18]. Recently, high-throughput mass spectrometry has greatly increased our ability to identify PTM sites [14, 22, 23, 20], increasing the number of known PTM sites from *<*1000 in 2002 to *>*440,000 today [12, 8]. Nonetheless, *>*65% of the motifs for the 518 human kinases remain uncharacterized [13].

In parallel with these experimental efforts is an ongoing stream of research devoted to characterizing PTM sites *in silico*. The PTM motif discovery problem differs from traditional protein sequence motif discovery in four important ways: (1) the candidate motif sites can be aligned relative to the (known) PTM, (2) the number of identified sites can be quite large, (3) the motifs tend to have low information content, and (4) the set of identified sites is usually contaminated with false positive identifications. Note that, like traditional motif discovery, the data set may contain a mixture of motifs, each corresponding to different modifications. The latter is particularly relevant to phosphorylation, where different types of kinases phosphorylate variant sequence motifs.

The current study offers three primary contributions. First, we provide a rigorous approach for estimating the statistical confidence associated with motifs discovered by the most widely used PTM motif discovery algorithm, motif-x. Our approach employs the Fisher exact test to measure the enrichment of the motif in the input peptides (the “foreground peptides”) relative to a control set of peptides (the “background peptides”). This methodology has been used previously by (non-PTM) motif discovery tools such as MEME [1] and DRIM [5]. We show that the motif *p*-values our approach reports accurately estimate the probability of discovering a motif at least as discriminative as the given one in shuffled versions of the foreground peptides.

Second, we describe a new software tool, MoMo (for “<u>Mo</u>dification <u>Mo</u>tifs”), that allows MEME Suite users to identify protein PTM motifs. MoMo fully reimplements the two most widely cited PTM motif discovery tools—motif-x [19] and MoDL [17]—and enhances them in several ways (Table 1). MoMo is fully integrated into the MEME Suite of motif-based sequence analysis tools, and is the only PTM motif discovery tool currently available that provides both a web server for interactive use as well as downloadable source code. In addition to implementing the novel statistical confidence estimation procedures described above, MoMo offers a number of improvements, as follows:

- MoMo’s implementation of motif-x uses a more accurate approach to calculate binomial *p*-values, which allows consistent motif discovery in much larger input datasets.
- MoMo web site users can access the proteomes of over 4000 organisms, compared with only eight organisms currently supported by the motif-x web server.
- MoMo users can upload larger proteome files than is possible using the motif-x webserver (80 megabytes vs 25 megabytes), and MoMo jobs are limited to 180 minutes run time, rather than motif-x’s limit of only 15 minutes.
- Unique among PTM motif discovery algorithms, MoMo can, in a single run, discover separate motifs for each combination of amino acid residue and modification mass (e.g., for phosphorylation, three separate motifs for serine, threonine or tyrosine), or MoMo can report a single motif for each modification mass, combining all peptides with that modification mass regardless of the modified residue.
- Another unique feature is MoMo’s ability to directly read peptide spectrum match (PSM) output formats from popular mass spectrometry search engines, such as Comet, MS-GF+, Tide, and Percolator (reviewed by Verheggen *et al.* [21]). For convenience, MoMo can directly filter PSM format files on any specified numeric field. MoMo can also extend the peptides in the PSM file to a given width using a user-specified proteome.

These enhancements make MoMo a flexible, easy-to-use and fully-featured solution for PTM motif discovery.

Third, in the course of our investigations, we uncovered several significant problems with the way PTM motifs are currently discovered using motif-x. We show that the common approach of using peptides drawn from the full proteome of the organism of interest as the background set often results in motifs that reflect only the different residue composition of the foreground peptides. These motifs may be incapable of distinguishing the foreground peptides from shuffled versions of those same peptides. Since motif-x motifs are supposed to be position-specific, we consider such motifs not to be statistically significant. We also demonstrate a disconnect between the “motif score” reported by the original version of motif-x and the statistical significance of the corresponding motifs. These observations demonstrate that the confidence estimation procedures we propose are needed in practice, and suggest that many of the hundreds of papers that cite motif-x may be reporting motifs that lack statistical support.

## 2 Methods

MoMo consists of a command line tool and a web interface. The command-line tool, henceforth referred to as “MoMo,” performs motif discovery in peptide sequences using one of three algorithms: motif-x, MoDL or simple alignment. We reimplemented the motif-x and MoDL algorithms completely from scratch from their published descriptions, and we verified that our versions behave identically with the web server and source code versions of motif-x and MoDL, respectively. The simple alignment algorithm is provided for convenience, and produces a single motif from all the sequences in the input that satisfy the criteria specified by the user (such as filtering on a numeric field in the PSM-formatted input). In the process of reimplementing motif-x, we enhanced it by adding the estimation of the statistical significance of the motifs it reports, by correcting an inconsistency in its algorithm, and adding the option to create the background peptides by shuffling the foreground peptides. In what follows, we will refer to MoMo’s enhanced version of motif-x as “motif-x*”.

### 2.1 MoMo command-line tool

The primary inputs to MoMo are one or more PTM files and a protein database file. The user can also specify the desired motif width, whether to remove modified peptides with ‘X’ (unknown residue) characters, whether to remove duplicate modified peptides, whether to output a single motif per modification mass or to separate isobaric PTMs by amino acid, and the minimum number of modified peptides required for a motif. MoMo produces as output an HTML report listing each discovered motif with its statistics, a sequence logo (in PNG format), and a list of the instances (modified peptides in its input) for each motif it discovers. In addition, MoMo summarizes its results in a tab-separated value (TSV) file, suitable for use with spreadsheet programs, and in a file of motifs in MEME motif format, which is suitable for use with the motif scanning (MAST and FIMO), enrichment (AME and CentriMo) and comparison (Tomtom) programs in the MEME Suite.

The PTM file input by the user to MoMo contain the modified peptide sequences. This file can be a set of fixed-width peptide sequences centered around the site of modification or a tab-delimited peptide-spectrum match (PSM) file. MoMo attempts to guess the format of the PTM input file. Fixed-width PTM files can be in either prealigned format, where each sequence is separated by a newline, or in FASTA format. With PSM format files, the user may specify the name of the column that contains the modified peptides, or they may specify one of the PSM program names listed in Table 2. As long as the modified peptides conform to one of the formats given in the last column of Table 2, then any tab-delimited file may be used. The user may also specify the name of another column in the PSM file that contains numeric data on which to filter the peptides, along with an (in)equality test (*<, ≤*, =, *≥,* or *>*) and a numeric threshold. Only peptides passing the specified filter will be included in MoMo’s analysis.

**Table 1:**
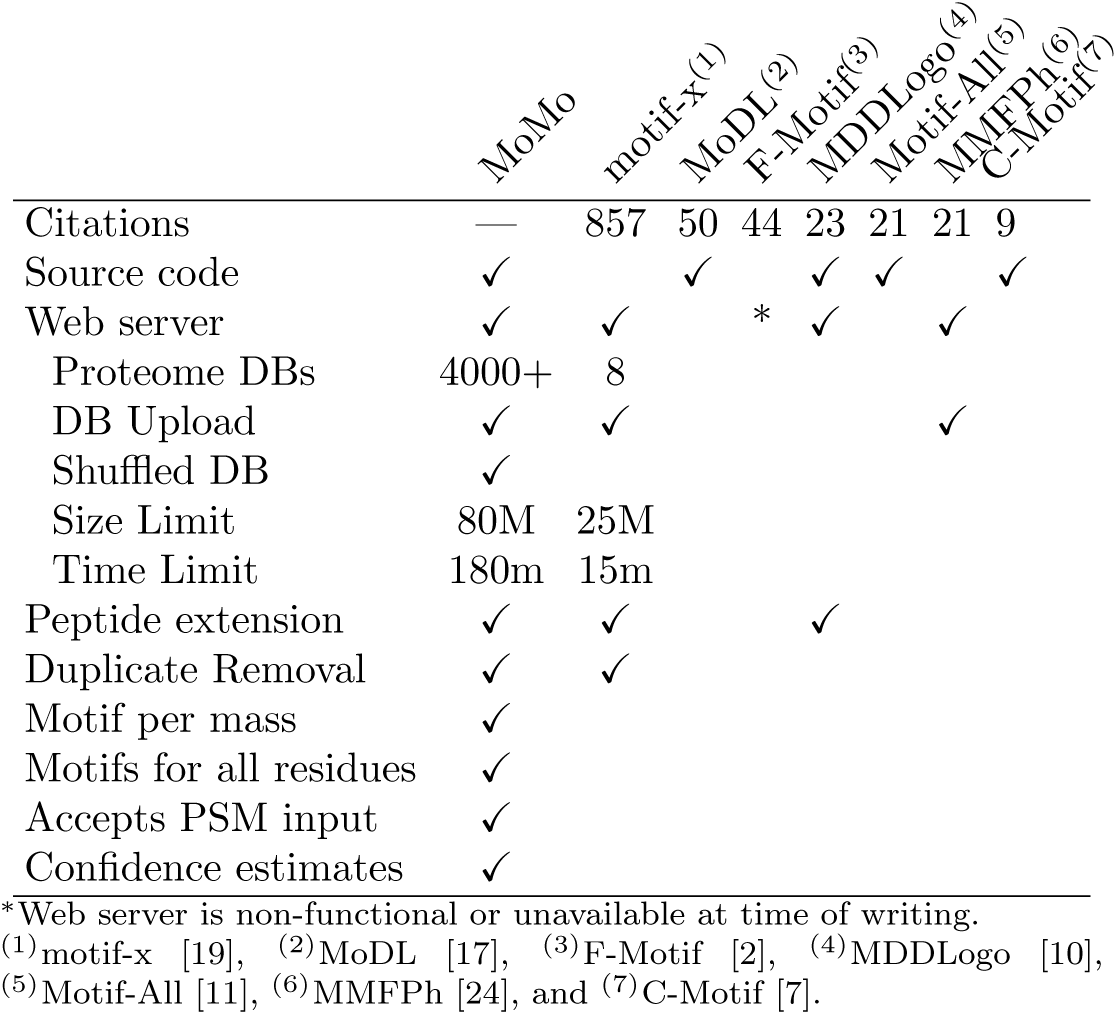
Comparison of PTM motif discovery tools.

**Table 2:**
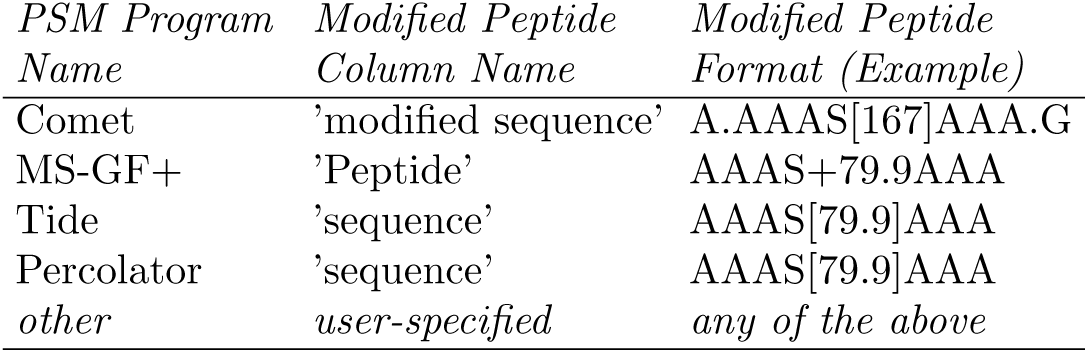
Peptide-spectrum match formats supported by MoMo.

| *PSM Program Name* | *Modified Peptide Column Name* | *Modified Peptide Format (Example)* |
| --- | --- | --- |
| Comet | ‘modified sequence’ | A.AAAS[167]AAA.G |
| MS-GF+ | ‘Peptide’ | AAAS+79.9AAA |
| Tide | ‘sequence’ | AAAS[79.9]AAA |
| Percolator | ‘sequence’ | AAAS[79.9]AAA |
| *other* | *user-specified* | *any of the above* |

MoMo’s other primary input is a protein database in FASTA format that typically contains the proteome of the organism from which the modified peptide data was gathered. MoMo uses the protein database to identify the context of each of the modified peptides in order to extend them on either or both sides to reach the requested motif width. MoMo does this by searching the protein database for matches to each modified peptide in its input and using the discovered context to provide flanking characters. If the discovered context overhangs one end of a protein sequence, the peptide is padded with the ‘X’ character. After this processing, all the modified peptides will have the requested motif width and be centered on a modified residue. MoMo allows the user to specify that any processed peptides that contain an unknown residue (’X’ character) at this point be eliminated from further processing. The user can also request that any processed peptides that are identical to another processed peptide in their central *N* residues be eliminated. (*N* must be shorter than the motif width.) The final set of fixed-width peptide sequences is referred to as the “foreground peptides.”

By default, MoMo looks for motifs where the central residue is the same for all instances of the motif. Unlike the original motif-x algorithm, MoMo does not require the user to specify the identity of the central residue, but returns motifs for all modified residues. Also, because each of the PSM formats supported by MoMo indicates the modification mass of each modified residue, the user can request that MoMo discover motifs where the central residue may vary, but the modification mass is fixed.

MoMo creates a set of “background peptides” for each of the central residues present in the foreground peptides. By default, for each central residue, the set of background peptides consists of a shuffled version of each of the foreground peptides. Shuffling of each peptide is done without disturbing the central residue. This mode of background creation is an enhancement to the original motif-x and MoDL algorithms, which only provide options to use an existing (unshuffled) proteome or to upload a user-created background file. With MoMo, if the user provides a background database such as the proteome of the organism providing the foreground peptides, then that database will be used in two ways. First, the database will be used to fill in any missing (’X’) residues in the foreground peptides and to extend the foreground peptides to the desired motif width. Second, the user may instruct MoMo to create the sets of background peptides it requires (one for each central residue in the foreground peptides) by extracting all contiguous peptides of the required length that contain the given central residue from the background peptides. MoMo then uses the background peptides to create the background model needed by motif-x or MoDL, and for computing motif *p*-values, as discussed below.

MoMo’s enhanced re-implementations of motif-x and MoDL are both written in C. The source code for motif-x was not made available by its authors, preventing it from being used in stand-alone mode. The original version of MoDL is written in MATLAB, and to run it requires a (paid) license for MathWorks. In contrast, MoMo is integrated into the MEME Suite, which is freely available as source code for non-profit use.

The original version of motif-x does not accurately calculate its objective function—the cumulative binomial probability— when its value is less than 10^-16^. This is due to a limitation of the `pbinom` function from the Perl module `Math::CDF` used by motif-x, as was previously noted by Chen *et al.* [2]. Our enhanced version of motif-x uses instead the regularized beta function,

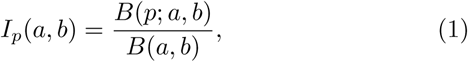

where *B*(*a, b*) is the beta function and *B*(*p*; *a, b*) is the incomplete beta function, to calculate the probability of *s* or more successes in *n* Bernoulli trials with prior probability *p* of success as

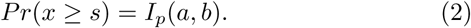

Our implementation of this function can accurately compute the cumulative binomial probability down to at least 10^−300^.

We enhanced motif-x by adding the estimation of the statistical significance of each reported motif. We follow the the practice of many existing motif discovery algorithms (e.g., MEME [1] and DRIM [5]) and use the Fisher exact test on the enrichment of the motif in the foreground peptides relative to the background peptides. Specifically, for each motif that motif-x* discovers, it calculates the *p*-value as the probability of *k* or more successes in *n* total draws from a population of size *N* with *K* total successes, where *k* is the number of foreground peptides with the motif, *K* is the number of foreground peptides, *n*-*k* is the number of background peptides with the motif and *N* -*K* is the number of background peptides. This probability is given by the hypergeometric distribution,

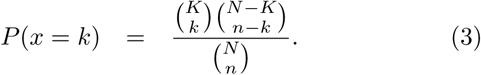

In addition to the Fisher exact test *p*-value, MoMo reports an adjusted *p*-value that takes into account the multiple testing inherent in the search performed by motif-x*. If motif-x* applied its binomial test *n* times in selecting the reported motif, then the adjusted *p*-value reported by motif-x* is *p*_*a*_=1 *-* (1 − *p*_*u*_)^***n***^, where *p*_*u*_ is the unadjusted *p*-value.

### 2.2 MoMo web interface

The MoMo web interface exposes the complete functionality of the command-line version of MoMo. The web input form allows the user to select the motif discovery algorithm to be used (motif-x, MoDL or simple alignment). The user can provide the modified peptides in PSM, FASTA or Raw format, and can provide multiple files that will be combined and analyzed as though they were a single file. The user can (optionally) upload a protein database, cut-and-paste a protein database, or select the proteome of one of the over 4000 organisms supported by the MEME Suite website. Additional options are provided for selecting the motif width and number of occurrences required, removing peptides with the ‘X’ character, eliminating duplicate peptides, creating a single motif per modification mass, and extracting the background peptides from the protein database. Algorithm-specific options are provided for the various thresholds and iteration limits required by the the motif-x and MoDL algorithms. Results are stored online, and the user is optionally notified of their availability via email.

### 2.3 Testing methodology and data

We used modified *Plasmodium falciparum* peptide data from Pease *et al.* [15] Supplemental Data 2 (their supplemental file `pr8b00062_si_003.xlsx`) and Supplemental Data 3 (their supplemental file `pr8b00062_si_004.xlsx`). As the protein database, we used the Ensembl version 38 *Plasmodium falciparum* proteome.

To test the accuracy of the *p*-values reported by motif-x*, we examined whether they follow the expected uniform distribution when the input peptides are random. We sorted together *p*-values of all the motifs reported by motif-x* when run with 10,000 shuffled versions of the Pease *et al.* [15] Supplemental Data 2 dataset, and we plotted the reported *p*-value as a function of its rank in the sorted list (Fig. 1). We ran motif-x* with the parameters `–width 13 –score-threshold 0.001 –protein-database` `E38_pfal.faa`, where `E38_pfal.faa` is the Ensembl version 38 *Plasmodium falciparum* proteome. Shuffled datasets for this experiment (and all other experiments reported here) were created by shuffling each peptide in the original dataset independently, conserving the central residue using the `fasta-shuffle-letters` tool provided with the MEME Suite. To compute the rank *p*-values shown in Fig. 1, we sorted the 10,000 reported *p*-values in decreasing order and computed the rank *p*-value of the *i*th reported *p*-value, *r*_*i*_, as *r*_*i*_= *i/*(*n* + 1), where *n*=10000. If the reported *p*-values are accurate, then the points (*r*_*i*_, *p*_*i*_) should lie along the line *y*=*x*.

**Figure 1:**
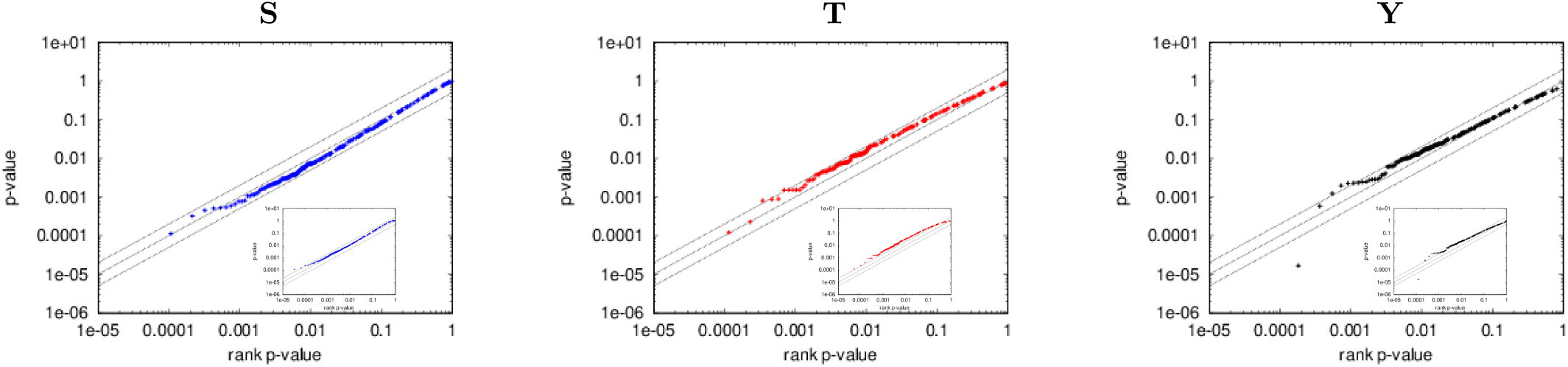
Accuracy of the *p*-values reported by motif-x^*\**^ in random data. The three panels show empirical assessments (Q-Q plots) of the statistical accuracy of the *p*-values reported by motif-x* for the motifs it discovers in 10,000 random datasets containing peptides centered on ‘S’, ‘T’ and ‘Y’ residues, respectively, when the *background peptides are shuffled versions of the foreground peptides*. Each panel shows results for motifs containing a given central residue. The main plot shows results for the *first* motif reported by MoMo, and the inset plot shows results for *all* motifs. Each point represents one motif reported by motif-x*, with *y* its *p*-value as reported by motif-x*, and *x* its rank *p*-value, *x* = 1*/*(*r*_*i*_ + 1)), where *r*_*i*_ is the rank of its *p*-value among those of the first (main panel) or all (inset panel) reported motifs. The three parallel lines show the curves (from top to bottom) for *y* = 2*x, y* = *x* and *y* = *x/*2, respectively. There are 3124, 518 and 233 peptides with central ‘S’, ‘T’ and ‘Y’ residues, respectively, in each of the 10,000 datasets.

To further test the accuracy of the *p*-values reported by motif-x*, we used the above results to empirically estimate the distribution of *p*-values reported by motif-x* on the original (unshuffled) Pease *et al.* [15] Supplemental Data 2 dataset. We then ran motif-x* on the *unshuffled* peptides in the Pease *et al.* [15] Supplemental Data 2 dataset, and we used the empirical distribution (estimated from motif-x* runs on the 10,000 shuffled versions of the dataset) to assign empirical *p*-values to each discovered motif. These results were used to construct the plots in Fig. 2, the table in Fig. 4A, and the plot in Fig. 4B.

**Figure 2:**
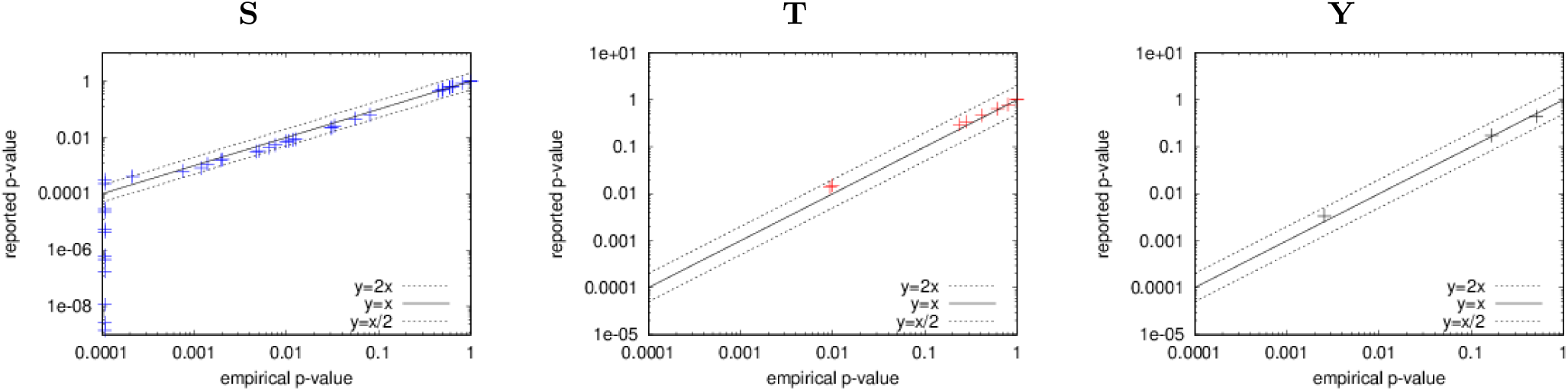
Accuracy of the *p*-values reported by motif-x^*\**^ in real data. The three panels show empirical assessments of the statistical accuracy of the *p*-values reported by motif-x* for the motifs it discovers in the Pease *et al.* [15] Supplemental Data 2 dataset when the background peptides are shuffled versions of the foreground peptides. Each point represents one motif reported by motif-x*, with *y* giving its *p*-value as reported by motif-x*, and *x* giving the *p*-value estimated empirically using 10,000 randomly shuffled versions of the same foreground peptides. There are 3124, 518 and 233 peptides with central ‘S’, ‘T’ and ‘Y’ residues, respectively, in the original input dataset, and in each of the 10,000 shuffled versions of it.

We repeated the above experiments using peptides extracted from the *Plasmodium falciparum* proteome by rerunning motif-x* with its `–db-background` switch. All other parameters and inputs remained the same. The output of these runs of motif-x* (on the original and 10,000 shuffled versions of the Pease *et al.* [15] Supplemental Data 2 dataset) were used to construct the empirical distribution, and to create the results shown in Fig. 3, Fig. 4C and Fig. 4D.

**Figure 3:**
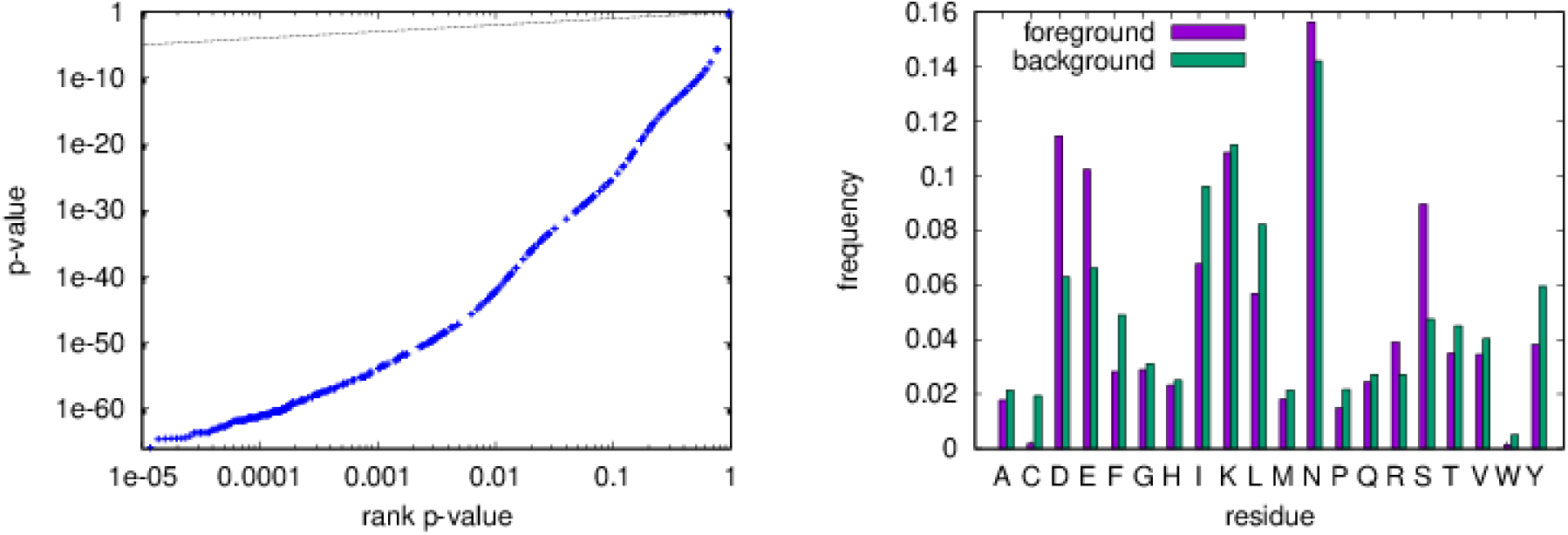
Bias of the *p*-values reported by motif-x^*\**^ when the foreground and background peptides have different residue frequencies. The left panel shows the empirical assessment (Q-Q plot) of the statistical accuracy of the *p*-values reported by motif-x* for 10,000 random datasets containing peptides centered on ‘S’ when the background peptides are extracted from a real proteome. Each point represents one motif reported by motif-x*, with *y* its *p*-value as reported by motif-x*, and *x* its rank *p*-value, *x* = 1*/*(*r*_*i*_ + 1)), where *r*_*i*_ is the rank of its *p*-value among all reported motifs. The right panel shows the residue distributions of the peptides in the foreground and background sets, excluding the central ‘S’ present in each peptide from the calculation. There are 3124 peptides with central ‘S’ in each of the 10,000 datasets.

**Figure 4:**
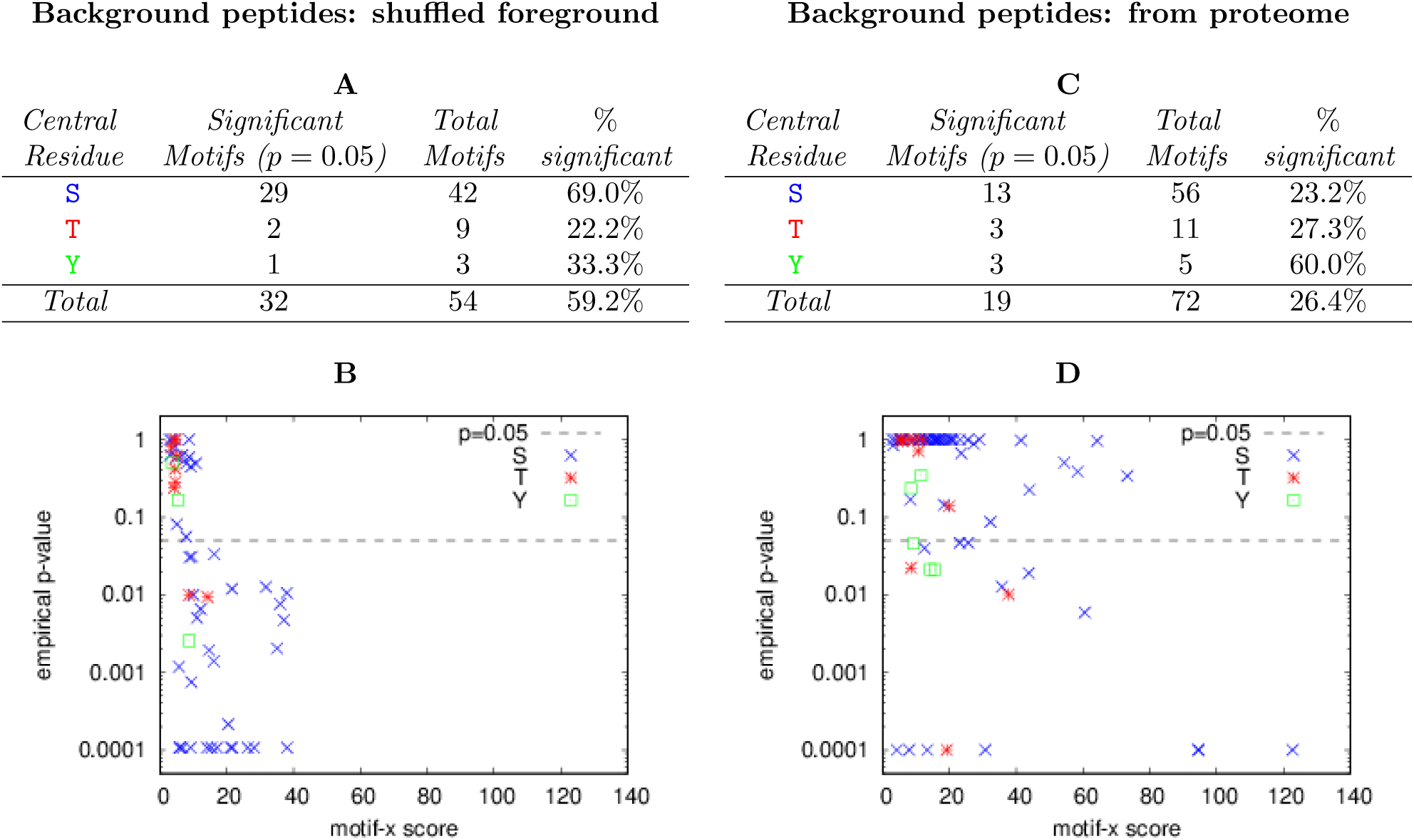
High motif-x scores are not indicative of high statistical significance. Panels A and B show the number of significant motifs reported by motif-x* and a scatter plot of motif significance vs. the reported motif-x score, respectively, when motif-x* uses shuffled foreground peptides as the background peptides. Panels C and D give the same information when motif-x* extracts the background peptides from the *Plasmodium falciparum* proteome. In both cases, the foreground peptides (input dataset) are from the Pease *et al.* [15] Supplemental Data 2 file. Empirical *p*-values are estimated from 10,000 runs of motif-x* on shuffled versions of the input dataset.

## 3 Results

### 3.1 MoMo’s enhanced version of motif-x accurately reproduces the published version

We checked that MoMo’s enhanced version of motif-x (motif-x*) produces results identical to those of the published version of motif-x, which is available only through an on-line web server. First, we reproduced the motif-x results from Fig. 3C in Pease *et al.* [15]. Running motif-x* with the same input files (the 196 sequences from Pease *et al.* [15] Supplemental Data 3 that are centered on ‘S’ and do not contain an ‘x’; the *Plasmodium falciparum* proteome as background) and motif-x* parameters equivalent to what Pease *et al.* used with motif-x (`–width 13 –score-threshold 0.00001 –min-occurrences 10 –db-background`), motif-x* outputs exactly the same three reported motifs.

Next, as a further check, we ran both versions of motif-x on the 3875 peptides in the Supplemental Data 2 file from Pease *et al.* [15]. The results produced by motif-x* are slightly different from the reported results, unless we supply the `–harvard` switch to motif-x*. With this switch, motif-x* still calculates binomial *p*-values for residue-position pairs correctly, but if any *p*-values are less than 10^***-*^16^**^, then it follows the practice of the original motif-x algorithm and chooses the residue-position pair among them that has the highest frequency. With the `–harvard` switch, motif-x* reports exactly the same 24 ‘S’ motifs as the published version of motif-x. Without the MoMo `–harvard` switch, motif-x* reports only 22 ‘S’ motifs on this set of peptides. Motifs `KxxxxxS*D, S*DD, S*DxxxxE` and `S*ND` are missing, and motifs `S*DxD` and `S*DxxD` are added. (We follow the motif naming convention in Pease *et al.* [15] of placing a ‘*’ after the modified residue, and indicating “don’t care” positions with the lower-case ‘x’ character.)

### 3.2 MoMo’s enhanced version of MoDL accurately reproduces the published version

We checked that MoMo’s enhanced version of MoDL (MoDL*) produces results identical to those of the published version of MoDL, which is available only from the command prompt within MATLAB. When run on the example input data provided with MoDL, MoDL* produced exactly the same motif as MoDL (DxxY), and the sequence of steps and description lengths computed at each step by MoDL* were identical with those computed by MoDL (to 6 decimal places). For this test, we ran MoMo with the switches `modl –width 13 –eliminate-repeats 0 –db-background`.

Note that, prior to completing the above test, we first had to make several corrections to MoDL. First, we corrected a bug in the MATLAB code that produces an error when MoDL fails to find a motif that improves the description length. In addition, we removed many bogus “peptides” from the sample background files distributed with the MoDL software, such as `CARBOXYLESTER` and `CHARACTERIZED`. Presumably, these sequences represent contamination from documentation inadvertently introduced when the MoDL authors created their example input files.

As a further check that MoDL* faithfully reproduces the published, MATLAB version of MoDL, we ran both versions of MoDL on the 3875 peptides in the Supplemental Data 2 file from Pease *et al.* [15]. Because MoDL requires that all the background peptides be of the same length and centered on the same residue, we extracted background files from the *Plasmodium falciparum* proteome, centered on each of the letters ‘S’, ‘T’ and ‘Y’, respectively. With each of these background files, MoDL* discovered the same motifs via the same sequence of steps, with the same description lengths (to 6 decimal places), as MoDL. The two ‘`S`’ motifs—`S*x[DE]` and `S*[DENPSV]x[DE]`—differ substantially from each of the 24 motifs discovered by motif-x* with the same input. The two ‘Y’ motifs found by MoDL* (`[DS]Y*` and `Y*[ES]`) are different from, but similar to, the five motifs found by motif-x* (`Y*S, SY*, Y*xS, SxY*` and `SxxxY*`). The three ‘`T`’ motifs found by MoDL* (`T*D, T*xxE` and `T*xD`) are vastly different from the eight motifs found by motif-x* (`T*DxE, T*D, T*xxE, ST*, T*xE, T*P, T*xD` and `T*xS`). As intended, MoDL* returns exactly the same results for all three central residues when run using the entire *Plasmodium falciparum* proteome as the protein database as it does when run separately on the extracted background files required by MoDL. Thus, MoMo’s enhanced version of MoDL gives the same results, but is much more convenient to use.

### 3.3 Computing the significance of motif-x motifs

We verified the accuracy of the motif significance estimates reported by motif-x* in two ways.

First, using sets of random peptides as input, we checked that the *p*-values followed the expected uniform distribution. We generated random peptides by shuffling real modified peptides, as described in Methods. The *p*-values of the *first* motif reported by motif-x* for each of the central residues contained in the random datasets are very uniform, as evidenced by the fact that the points main panels in Fig. 1 all lie close to the line *y*=*x*. When we consider *all* motifs reported by motif-x*, then the reported *p*-values tend to lie slightly above the line *y*=*x*, indicating that motif-x* may be slightly underestimating the significance of some of the motifs it reports. The reported *p*-values are therefore slightly conservative, and are generally within a factor of 2 of the true *p*-value of the motif.

Second, we measured the accuracy of the *p*-values reported by motif-x* when it is run on a real data set of modified peptides. In this case, we measured the accuracy of the *p*-values reported by motif-x* by comparing them to empirical *p*-values derived using Monte Carlo estimation (see Methods). The high accuracy of the *p*-values reported by motif-x* on the real peptide data is evident from the fact that the points (*p*_*e*_, *p*_*u*_) in Fig. 2 lie very close to the line *y*=*x*, where *p*_*e*_ is the empirical *p*-value, and *p*_*a*_ is the reported *p*-value. In the case of motifs with ‘S’ as the central residue, some of the motifs reported by motif-x* in the real dataset have *p*_*a*_ less than 1/10,000, the smallest empirical *p*-value possible in this experiment (points along the y-axis). We cannot definitively say that *p*-values below 10^***-*^4^**^ are as accurate as larger ones, but we can assert that those motifs are statistically significant, since none of the 10,000 random datasets produced a motif with a *p*_*a*_ as small.

### 3.4 Deriving the motif-x background from the proteome can be misleading

The above two evaluations used motif-x* in its default mode, where the program creates a set of background peptides by shuffling each of the foreground peptides, conserving its central residue. However, the motif-x algorithm is commonly run using a background derived from all suitable (unshuffled) peptides in the proteome. We hypothesized that searching for motifs in this fashion may lead to inaccurate *p*-values. The reason for the potential inaccuracy is that motif-x* may find a motif that discriminates even random foreground peptides from the background peptides if the two sets of peptides have very different residue distributions. This is obviously true in the extreme case where the foreground peptides are highly enriched for a single residue that is completely absent from the background peptides. In this case, any motif containing the residue will tend to be a good discriminator.

To test this hypothesis, we ran motif-x* on *random* data created by shuffling the peptides in a real dataset, and having motif-x* create background peptides from the corresponding proteome (Fig. 3). Specifically, the foreground peptides are shuffled versions of the 3124 peptides with a central ‘S’ residue in the Pease *et al.* [15] Supplemental Data 2 dataset. Motif-x* was run using its `–db-background` option and the Ensembl version 38 *Plasmodium falciparum* proteome. The motifs found by motif-x* in the shuffled foreground peptides tend to have extremely low reported *p*-values, rather than following the uniform distribution as they should (dotted black line in left panel of Fig. 3). This trend is due to motif-x* discovering motifs that contain *residues* that are highly enriched in the foreground peptides relative to the background peptides due to the difference in overall residue composition of the peptides in the Pease *et al.* [15] Supplemental Data 2 dataset compared to the proteome (right panel of Fig. 3). The residues ‘`D`’, ‘`E`’ or ‘`S`’ are relatively enriched in the foreground peptides, and, consequently, in shuffled versions of it, so motifs containing them tend to be discovered by motif-x* (and motif-x), and have high discriminative power. However, the biological significance of the `D`-, `E`- or `S`-rich motifs found in shuffled input datasets is questionable (to say the least), and suggests that the failure of the motif-x algorithm to adjust for the relative residue composition of the foreground and background peptide sets may be a major weakness. Accordingly, if the user opts to employ an unshuffled background via the `–db-background`, motif-x* includes a warning in its output about the potential for inaccurate *p*-values.

### 3.5 Not all motifs reported by motif-x are significant

Although the two publications describing the motif-x algorithm claim that the motifs it reports are statistically significant, we found that this is not the case using our definition of significance (see previous section). Likewise, we found that the motif scores reported by motif-x do not seem to be a good guide for judging the significance of motifs. Our conclusions about the lack of significance of many motif-x motifs, as well as the lack of correlation between motif score and motif significance apply regardless of whether shuffled foreground peptides or peptides extracted from the proteome are used as background peptides by motif-x.

The original motif-x paper states [19]:

> “Despite the statistical significance of every motif extracted, heuristic scores for the motifs were calculated as the sum of the negative log of the binomial probabilities used to generate the motifs

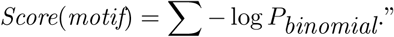

A later paper by the developer of motif-x describing the protocol for using motif-x states [3]:

> “The ‘motif score’ is calculated by taking the sum of the negative log probabilities used to fix each position of the motif. As such, higher motif scores typically correspond to motifs that are more statistically significant as well as more specific (i.e., greater number of fixed positions).”

However, we found that many motifs reported by motif-x* (and, hence, by motif-x) are *not* statistically significant, and that the motif scores it reports are *not* highly correlated with motif significance. For example, of the 54 motifs found by motif-x* in the 3875 peptides in Supplemental Data 2 from Pease *et al.* [15] using the (default) shuffled foreground peptides as background peptides (Fig. 4A), only 32 (59.2%) are statistically significant (*p*=0.05), as estimated using the Monte Carlo simulation. If background peptides are extracted from the *Plasmodium falciparum* proteome (Fig. 4C), then only 26.4% of the motifs reported by motif-x* are significant. In other words, in these two examples, about 40% and 75%, respectively, of the reported motifs are *not* enriched in the foreground peptides compared to the background peptides at the *p*=0.05 level, as measured by the Fisher exact test. Furthermore, especially when the background peptides come from the proteome (Fig. 4D), the motif score reported by motif-x* is not strongly correlated with motif significance. Many non-significant motifs (points above the *p*=0.05 line in Fig. 4D) have motif scores larger than motifs with empirical *p*-values at or below 0.0001. Conversely, when shuffled versions of the foreground peptides are used as the background peptides, some highly significant motifs reported by motif-x* have motif scores close to zero, and less than those of some non-significant motifs.

As a second example, we ran motif-x* on the much smaller dataset (196 peptides) used by Pease *et al.* [15] (their Supplemental Data 3 file), and used peptides from the *Plasmodium falciparum* proteome as the background, to reproduce their Fig. 3C, as discussed at the start of the Results section. As noted above, this analysis yields the same three, previously reported motifs. However, we find that only one of the three motifs is statistically significant at the *p*=0.05 level when we compute their empirical *p*-values by running motif-x* on 10,000 shuffled versions of the same dataset (Fig. 5A). Based on this analysis, the `RxxS*` motif discovered by motif-x, and which Pease *et al.* [15] claim is a known phosphorylation target for the AGC-kinases, is *not* significantly enriched in the foreground sequences relative to the background sequences. On the other hand, the `S*xD` motif discovered by motif-x is significantly enriched at the *p*=0.05 level (Fig. 5A) but is not a known kinase target. As a further check, we also ran motif-x* using shuffled foreground peptides as the background (its default mode), and note that the program still finds the `S*xD` motif, but that it is not significant at the *p*=0.05 level (Fig. 5B). The `RxxS*` motif discussed by Pease *et al.* [15] is not found at all in this setting. These results suggest that the relevance of the motif analysis reported in Pease *et al.* [15] may be low.

**Figure 5:**
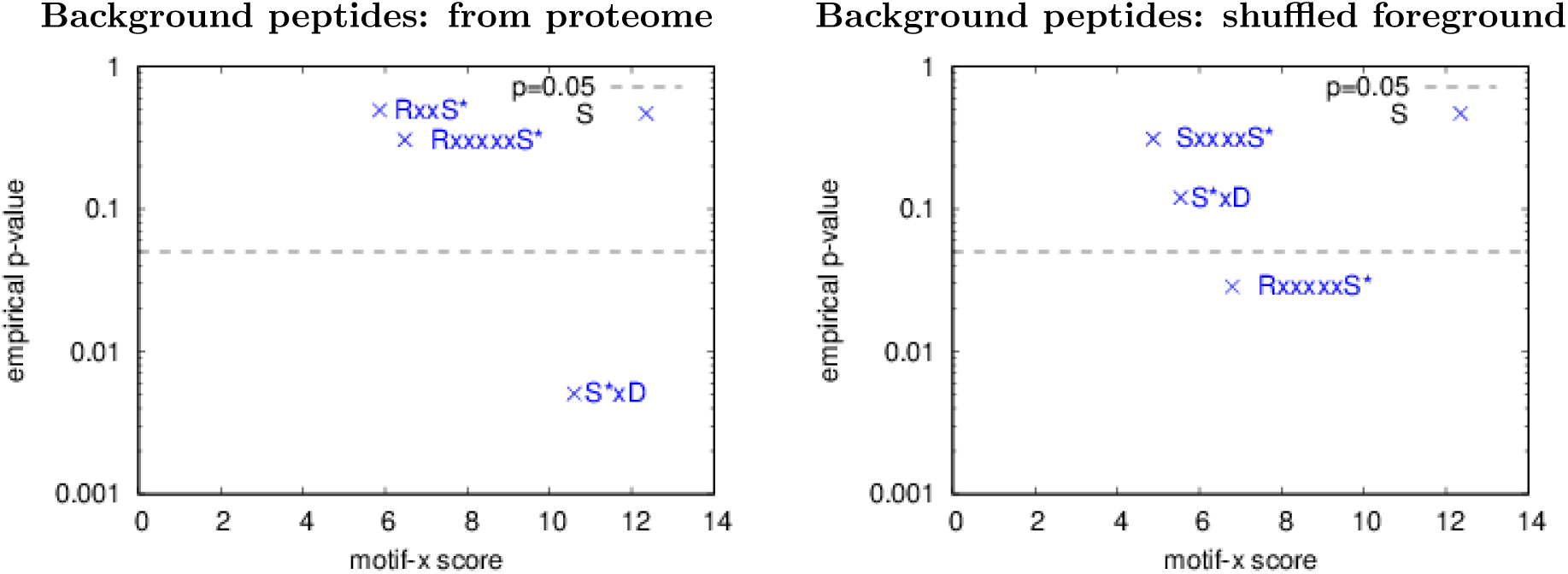
Only one of the three motif-x motifs reported in Pease *et al.* [15] is statistically significant. The panels show the motif-x score and empirical *p*-values of the motifs found by motif-x* using the peptides in Supplemental Data 3 and the Ensembl version 38 *Plasmodium falciparum* proteome, and peptides from the proteome (panel A) or shuffled foreground peptides (panel B) as the background peptides. The minimum number of occurrences parameter is 10 in both panels, and the minimum motif-x score parameter is 0.00001 in panel A, and 0.0001 in panel B. (Only the S*xD motif is found if the minimum score is set to 0.00001 in panel B.)

## 4. Discussion

We have shown that MoMo’s version of motif-x reports accurate motif *p*-values when run in its default mode, where the background peptides for motif-x are created by MoMo by shuffling the foreground (input) peptides. This provides users of MoMo with an accurate estimate of the statistical significance of the motifs its version of motif-x reports. This is important, because our results show that, contrary to what has been suggested [19, 3], there is no guarantee that the motifs discovered by motif-x are statistically significant.

Another important point raised by our study is that, for PTM motif discovery, the ideal background set may not be all the peptides in the proteome that have a given residue in their center. Often, the set of modified peptides in which we wish to find motifs has a very different residue distribution than the proteome as a whole. As a result, motif discovery algorithms (like motif-x) may find motifs that are very good at distinguishing the foreground peptides from the background peptides, but do so only because the motifs capture this difference in residue composition. This is illustrated by the fact that MoMo’s version of motif-x finds only 19 statistically significant motifs when using the proteome-based background, compared with 32 significant motifs using the shuffled input background (Fig. 4). In that experiment, only 26.4% of the motifs found by motif-x using the proteome-based background were significant at the *p*=0.05 level.

MoMo provides users with easy access to improved versions of motif-x and MoDL, and can provide accurate estimates of the statistical significance of motif-x motifs. Significance estimates are only accurate when MoMo is allowed to create the background dataset by shuffling the input peptides. With other backgrounds (e.g., proteome or user-created background sets), and with MoDL motifs, users should use other means to assess the statistical significance the PTM motifs reported by MoMo and other algorithms. One approach is to estimate the empirical *p*-values of motifs by running the motif discovery algorithm many times on shuffled versions of the input peptides. The MEME Suite, of which MoMo is a part, provides the `fasta-shuffle-letters` script, which can shuffle peptides, conserving the central residue.

